# High levels of antibiotic resistance gene expression among birds living in a wastewater treatment plant

**DOI:** 10.1101/462366

**Authors:** Vanessa R. Marcelino, Michelle Wille, Aeron C. Hurt, Daniel González-Acuña, Marcel Klaassen, John-Sebastian Eden, Mang Shi, Jonathan R. Iredell, Tania C. Sorrell, Edward C. Holmes

**Affiliations:** Marie Bashir Institute for Infectious Diseases and Biosecurity, Sydney Medical School, The University of Sydney, Sydney, NSW 2006, Australia.; Westmead Institute for Medical Research, Westmead, NSW 2145, Australia.; School of Life & Environmental Sciences, Charles Perkins Centre, The University of Sydney, Sydney, NSW 2006, Australia.; WHO Collaborating Centre for Reference and Research on Influenza, at The Peter Doherty Institute for Infection and Immunity, Melbourne, VIC 3000, Australia.; Laboratorio de Parásitos y Enfermedades de Fauna Silvestre, Facultad de Ciencias Veterinarias, Universidad de Concepción, Concepción 3349001, Chile.; Centre for Integrative Ecology, School of Life and Environmental Sciences, Deakin University, Geelong, VIC 3216, Australia.

**Author notes:** **Corresponding Author:** Vanessa R. Marcelino, Marie Bashir Institute for Infectious Diseases and Biosecurity, Sydney Medical School, The University of Sydney, Sydney, NSW 2006, Australia.

**Keywords:** Metatranscriptomics, Microbiome, Birds, Resistome, Antimicrobial resistance

## Abstract

Antibiotic resistance is rendering common bacterial infections untreatable. Wildlife can incorporate and disperse antibiotic resistant bacteria in the environment, such as water systems, which in turn serve as reservoirs of resistance genes for human pathogens. We used bulk RNA-sequencing (meta-transcriptomics) to assess the diversity and expression levels of functionally active resistance genes in the microbiome of birds with aquatic behavior. We sampled birds across a range of habitats, from penguins in Antarctica to ducks in a wastewater treatment plant in Australia. This revealed 81 antibiotic resistance genes in birds from all localities, including β-lactam, tetracycline and chloramphenicol resistance in Antarctica, and genes typically associated with multidrug resistance plasmids in areas with high human impact. Notably, birds feeding at a wastewater treatment plant carried the greatest resistance gene burden, suggesting that human waste, even if it undergoes treatment, contributes to the spread of antibiotic resistance genes to the wild. Differences in resistance gene burden also reflected the birds’ ecology, taxonomic group and microbial functioning. Ducks, which feed by dabbling, carried a higher abundance and diversity of resistance genes than turnstones, avocets and penguins, that usually prey on more pristine waters. In sum, this study helps to reveal the complex factors explaining the distribution of resistance genes and their exchange routes between humans and wildlife.

## Introduction

Tons of antibiotics are used annually in clinical and agricultural settings worldwide. Food animals alone consumed over 130,000 tons of antibiotics in 2013 (1), and antibiotic usage by humans increased 65% between 2000 and 2015, reaching 34.8 billion defined daily doses (2). The resulting proliferation and spread of bacteria that are resistant to antibiotics poses a major health and economic threat (3). Genes for the production of antibiotics and antibiotic resistance determinants are naturally found in some microbial species and their presence in the environment is not necessarily an indication of human impact (4, 5). However, the use of antibiotics in clinical and agricultural settings selects for bacteria carrying resistance genes. When these genes are encoded in mobile elements, such as plasmids and conjugative transposons, they can be readily transmitted via horizontal gene transfer between environmental bacteria and clinically important pathogens (i.e. acquired resistance genes). Multiple resistance genes can be present in a single mobile element and the spread of plasmid-borne resistance has jeopardized the efficacy of many antibiotics, including β-lactam drugs of last resort (6, 7).

Both the environment and wildlife are major sources and reservoirs of resistance gene diversity (8, 9). The ecological niches and behavior of birds make them particularly likely to transport antibiotic resistant bacteria. Migrating bird species transport pathogens which may contain antibiotic resistance genes across large distances (10, 11). Birds also serve as sensitive bioindicators of environmental contamination with antibiotic resistant bacteria (10, 12-16). For instance, ESBL-producing *Escherichia coli* were found to occur over 3 times more frequently in gulls than in humans in the same region (15). Bacteria resistant to β-lactam and tetracycline drugs are commonly found in the gut microbiome of birds, especially in scavenging and aquatic species, such as waterfowl, gulls and waders (10, 12, 14, 17-20). Aquatic bird species likely acquire these genes through contact with contaminated water. Human sewage is enriched in antibiotic-resistant bacteria, which are only partially removed during the water treatment process (21-26). Birds in contact with wastewater treatment influents or effluents could therefore be at increased risk of acquiring these genes, although empirical data to support this idea are scarce (8).

While the majority of studies on birds were based on *in vitro* assessments of bacterial cultures, the development of culture-independent sequencing techniques has substantially expanded our knowledge of the environmental reservoir of resistance genes (7, 9, 27-32).

Among these techniques, sequencing the entire set of transcribed (i.e. expressed) genes via ‘meta-transcriptomics’ has rarely been used in the context of antibiotic resistance, despite its advantages. In particular, use of meta-transcriptomics allows data to be obtained from the entire microbial population, with a focus on functionally active genes. This is important because genetic material is a metabolic burden and genes that are not essential tend to be lost (33-36). In the absence of antibiotics, it is likely that resistance genes are regularly lost by bacteria, either by large deletions or gradual deactivation (erosion). Other high-throughput techniques, such as DNA-based metagenomics, cannot distinguish recently deactivated resistance genes from their functional relatives. An alternative is to clone inserts from environmental strains into cultivable vectors (e.g. *E. coli*), select for resistance *in vitro* and then sequence their genomes (e.g. 17, 22). However, this approach can result in bias towards genes present in organisms closely related to the cloning vector (7). Meta-transcriptomics does not have this limitation as the transcripts of all microorganisms are assessed using bulk RNA sequencing. To our knowledge, only two studies have used meta-transcriptomics to report on the presence of resistance genes that are functionally active under natural conditions in human and environmental samples (37, 38).

We used meta-transcriptomics to assess the diversity and abundance of antibiotic resistance genes actively transcribed in the microbiome of water birds of Australia and penguins in Antarctica. Birds were sampled across a range of habitats, from remote places in Antarctica and Australia, beaches in Melbourne, the second largest city in Australia, to the ponds of a waste water treatment plant (WWTP) processing half of Melbourne’s sewage. We specifically tested whether ducks from the WWTP harbor a higher diversity and abundance of acquired resistance genes, as might be expected given their exposure to partially treated human waste. Additionally, we explore possible associations between resistance gene burden and intrinsic bird traits such as feeding behavior, taxonomic order and gut functional profile (expression of metabolic pathways by the microbiome).

## Results

Microbiome samples from 110 birds, grouped into 11 libraries (Table S1), contained transcripts corresponding to 81 unique antibiotic resistance genes, previously associated with phenotypic resistance to nine classes of antibiotics (Fig. 1, Table S2). These results only include acquired resistance genes, which are most commonly spread among bacteria via mobile genetic elements, and do not include resistance mediated by chromosomal mutations (e.g. in housekeeping genes). Resistance to tetracyclines and phenicols (chloramphenicol and florfenicol) was present in samples from all bird orders and in all locations, except for one site in Antarctica where phenicol resistance was not detected.

**Figure 1.**
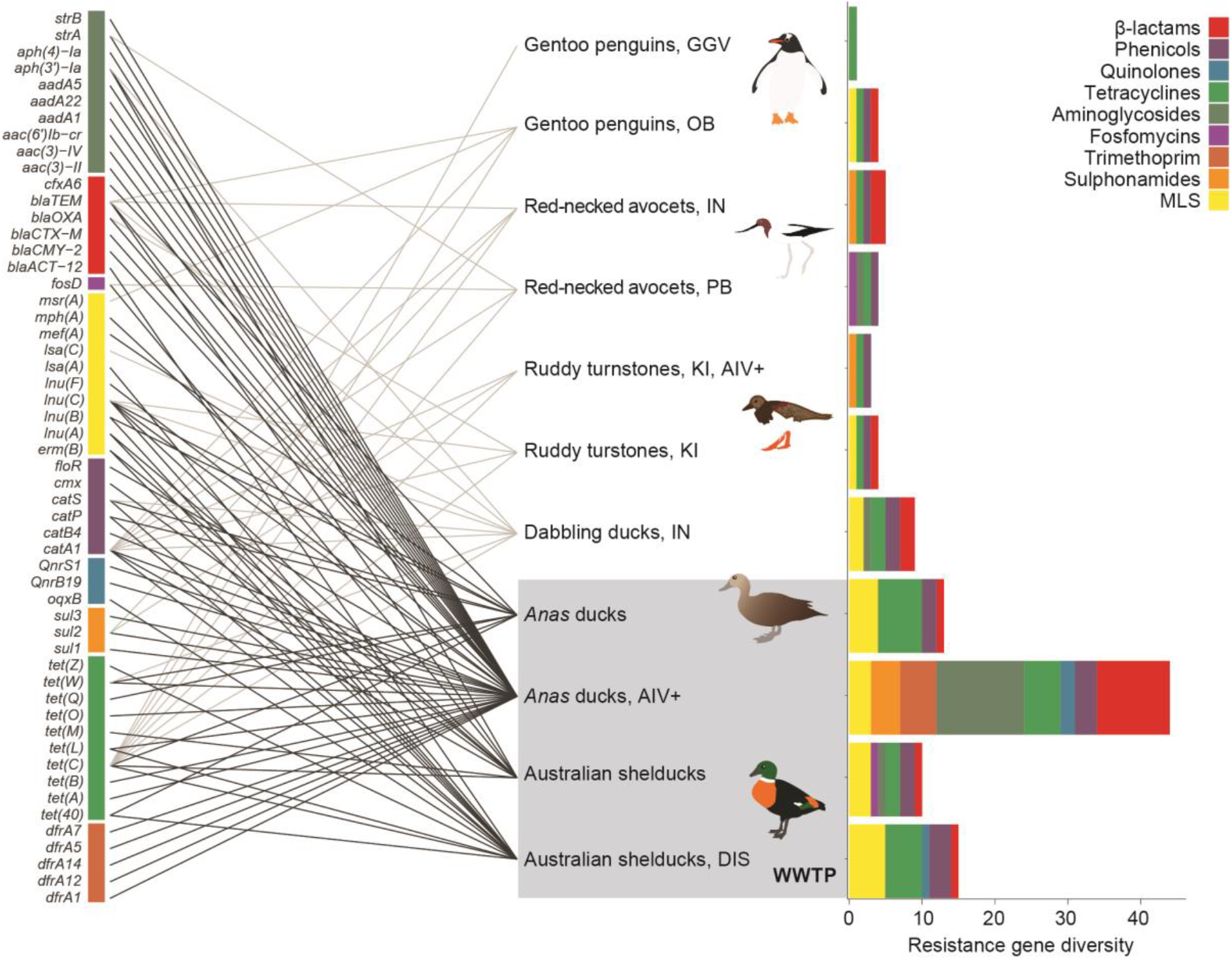
Antibiotic resistance genes expressed in the microbiome of wild birds. The graph on the right shows the diversity of resistance genes observed in each library (containing a pool of 10 individual birds each), colored by the drug class to which these genes confer resistance. Closely related gene variants were merged into one category (see Table S2) for representation on the left side of the figure. Lines link genes to the libraries where they were found, and dark lines indicate the genes observed in the Wastewater treatment plant (WWTP) in Melbourne, Australia. PB = Western Port Bay, Melbourne area, Australia; KI = King Island, Bass Strait, Australia; IN = Innamincka reserve, Australia; OB = O’Higgins Base, Antarctica; GGV = Gabriel González Videla Base, Antarctica. Libraries of birds infected with avian influenza virus are indicated with ‘AIV+’, and the library of diseased birds is indicated with ‘DIS’. MLS = Macrolides, Lincosamide and Streptogramin B resistance. Bird drawings: M. Wille.

### Anthropogenic impact

Birds foraging at the partially treated lagoons of a wastewater treatment plant (the last stage of the wastewater treatment process, after aerating and decanting has taken place) had a significantly higher diversity and abundance of antibiotic resistant genes, as well as a significantly higher number of antibiotic classes against which these genes confer resistance (Kruskal-Wallis *p*<0.05, Fig. 2). For simplicity, we refer to ‘resistance gene burden’ or ‘resistance load’ as the resistance gene diversity, abundance (i.e. gene expression levels) and number of antibiotic classes to which these genes confer resistance. Most notably, ducks (order Anseriformes) foraging at the WWTP harbored 86% of the resistance gene diversity observed, most of which occurred exclusively at the WWTP (Figs. 1, 2, Table S2).

**Figure 2.**
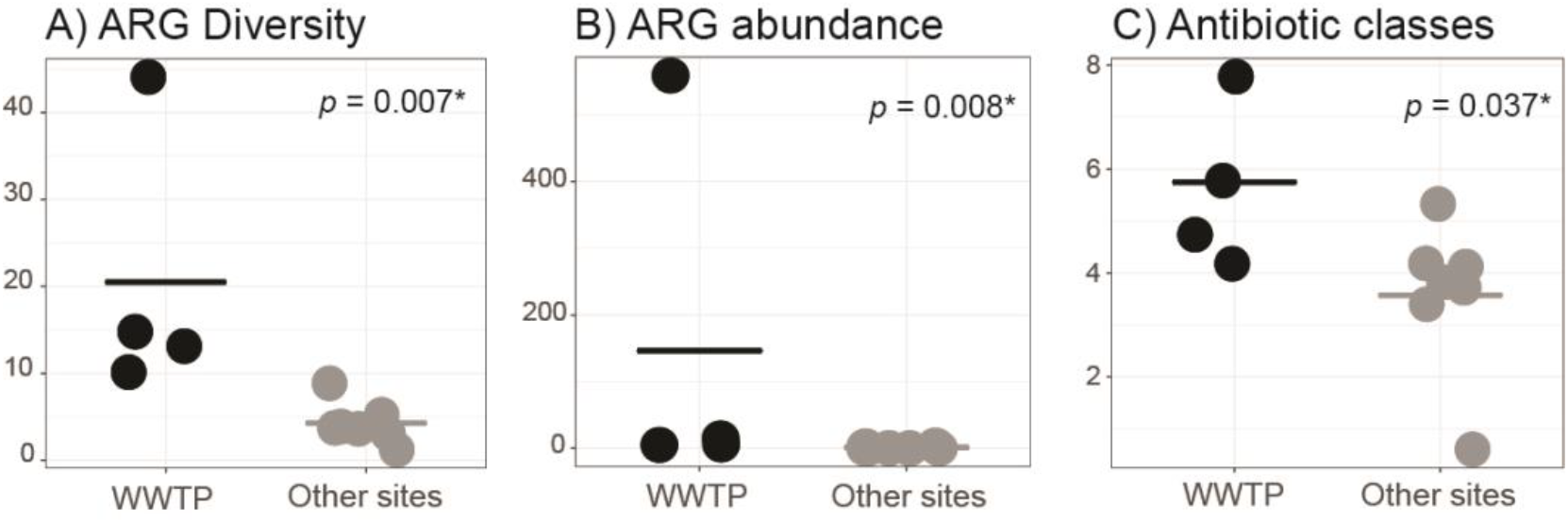
Diversity and abundance of antibiotic resistance genes (ARG) in birds foraging in a wastewater treatment plant (WWTP) compared with birds from other sites in Australia and Antarctica. Differences between groups were assessed with a Kruskal-Wallis test and *p*-values are given. Resistance gene abundances were estimated based on a stably expressed host gene.

When only ducks were considered in comparing the effects of wastewater, we observed that those from the WWTP carried more resistance genes than ducks from the remote Innamincka reserve, located in the interior of Australia (Fig. 1, Table S3): ducks from the Innamincka reserve carried nine resistance genes, fewer than the number observed in any library from the WWTP (average 20.5, +/-15.8 SD). The abundance of these genes was also smallest in ducks from Innamincka (2.9, compared with an average of 146.1, +/- 275.2 SD in ducks from the WWTP). The number of antibiotic classes to which these genes confer resistance did not differ substantially between sites (5 antibiotics in birds from Innamincka, compared with 5.7, +/- 1.7 in birds from the WWTP).

To take into account potential confounding variables, we also analyzed libraries by individual collection localities (Fig S1), and without including diseased birds or birds infected with avian influenza in the analyses (Fig. S2). The results consistently indicated that birds from the WWTP have a higher resistance gene burden than birds from other localities. Importantly, an additional PCR-based assessment of the resistance genes in individual birds from two libraries (n=20 samples) confirmed the results obtained using meta-transcriptomics: we observed 68 resistance gene occurrences (amplifications) in samples from the WWTP and 12 occurrences in other sites (Kruskal-wallis *p*=0.0023, Supplementary Materials, Fig S3).

Samples from gentoo penguins (*Pygoscelia papua*) collected in two localities next to research bases in Antarctica, contained five resistance genes in total, conferring resistance against β-lactams (*bla*_TEM_), tetracyclines (two variants of *tet*(C)), chloramphenicol (*catA1*) and erythromycin (*msr*(A)) (Table S2). The erythromycin-resistance gene, which confers resistance to Macrolides, Lincosamide and Streptogramin B, was observed in penguins only. Penguins living near the research base with the largest human population (O’Higgins Base) contained more antibiotic resistance genes (four genes - *bla*_TEM,_, *msr*(A), *catA1*, and *tet*(C)), than those living next to the more remote Gabriel González Videla Base (one *tet*(C) gene).

### Host traits and functional context

Our sampling design included birds from a range of habitats and species, which will impact their microbiome and possibly their propensity to carry antibiotic resistance genes. Shelducks and *Anas* ducks (Anseriiformes) feed by dabbling (filtering water). Turnstones and avocets (Charadriiformes) commonly prey on invertebrates, and penguins (Sphenisciformes) prey on fish. Dabbling ducks live in a range of habitats, including nutrient-rich and heavily altered environments. The majority of ducks analyzed here were sampled at the WWTP: 40 samples (4 libraries) at the WWTP and 10 samples (1 library) in a pristine site. Turnstones and penguins on the other hand inhabit pristine habitats. Host taxonomic order therefore serves as a proxy for the ecology of the birds analyzed here. Our results indicated that ducks contained the greatest diversity and abundance of resistance genes, while penguins contained the lowest resistance load (Fig. 1 and Fig. S4).

Host ecology is intrinsically linked to microbiome function. By investigating how microbiomes functionally differ among bird orders and collection sites we can gain insights into why some hosts harbor more resistance genes than others. We characterized the metabolic pathways expressed by the microbial community (that is, their functional profile, Table S4). Some of the metabolic pathways observed were produced by common human pathogens (e.g. *E. coli*), but a large proportion of the metabolic products (91%) could not be associated with particular bacterial genera (Table S4). Compared with the human gut, the microbiome of wild animals is far less characterized, and it is expected that several bacterial species remain undetected. Principal coordinate analyses showed that ducks (from Innamincka reserve and from the WWTP) have a distinct microbial metabolism (i.e. set of metabolic pathways) when compared with birds from other sites (Fig S5). We statistically assessed the distinctiveness of functional profiles between sites and bird orders using Random forest analysis, a machine learning approach based on classification trees that has a suitably high discriminating power for use in microbial ecology (39). This analysis revealed a clear distinction (zero out-of-bag classification error) in the functional profiles between birds from the WWTP and other sites, and between Anseriformes and the two other bird orders that comprised the data set (Charadriiformes and Sphenisciformes; Table S5).

The bird microbiome, and consequently its functional profile, can also be affected by pathogens (40). We sampled birds with avian influenza virus infection and Newcastle disease symptoms; potential associations between these infections and antibiotic resistance are discussed in the Supplementary Materials.

## Discussion

This study shows that clinically important and functional antibiotic resistance genes are widespread, even in birds from areas as remote as Antarctica, and that the resistance gene load is significantly higher in birds living in lagoons of a wastewater treatment facility. Although resistance genes can be found in natural environments regardless of human influence (4, 5), our results indicate that contact with human waste – even if it goes through sewage treatment – appears to have a strong impact on the acquisition of antibiotic resistance genes by avian wildlife.

The resistance genes observed here encompass the three major resistance mechanisms of relevance to human infection: (i) drug inactivation, (ii) reduced influx of antibiotics into bacterial cells or increased efflux from cells and (iii) alteration in, or overexpression of, the antibiotic target (7, 41). The observed resistance genes conferred resistance against nine classes of antibiotics (Fig. 1). This number is slightly higher than the six classes of antibiotic resistance observed in humans, pigs, sponges and environmental samples in another study using meta-transcriptomics (37). Among the most common were genes conferring resistance to β-lactam drugs, which is one of the oldest and most widely used antibiotic classes. Genes conferring resistance to aminoglycosides and tetracyclines were also common, in agreement with studies reporting these genes in human-impacted soils and sewage (22, 27, 29, 31).

Some of the resistance genes observed are particularly concerning for public health. *bla*_CTX-M_ genes, observed exclusively in birds from the WWTP, play a key role in widely disseminated and highly resistant *E. coli* and *Klebsiella pneumoniae* strains (42). A fosfomycin resistance gene (*fosD*) was found in birds from metropolitan Melbourne (WWTP and Western Port Bay). Fosfomycin was discovered over 40 years ago, it is uncommonly used in humans, but the low resistance levels against this drug have led to a renewed interest in its therapeutic use (43). One of the bird libraries from the WWTP contained a florfenicol resistance gene, which was first observed in *Salmonella typhimurium* (44). Florfenicol is restricted to livestock and veterinary use. It is possible that the presence of this gene is due to the administration of florfenicol to pets and wildlife within the WWTP catchment range. The florfenicol gene has been observed co-located with other resistance genes in integrons and plasmids (44, 45). It is therefore also possible that this gene is found in the WWTP due to co-selection with other genes. We also found resistance against chemically synthesized antibiotic classes, such as quinolones and sulphonamides, which are not expected to be widespread in the environment (unlike naturally produced antibiotics such as penicillin, which is derived from fungi). Quinolone drugs can persist in the environment for long periods (46) and despite being a synthetic drug, the origins of quinolone resistance were traced back to aquatic bacterial species (47). Therefore it is perhaps unsurprising that these genes are found in birds with aquatic behavior (also reported in 19). It is noteworthy, however, that quinolone resistance was only observed in birds near the WWTP, suggesting that these genes most likely derive from bacteria of human origin. One of the WWTP libraries also contained the *aac(6)-Ib-cr* gene (100% identity with clinical isolates), which confers resistance to quinolones and aminoglycosides and is often localized in multidrug resistance plasmids. First reported in Shanghai in 2003, this gene has already been found in several parts of the world, including in a recent report of multidrug-resistant *Salmonella* in Australia (7, 48, 49).

The distinct ecological niche and microbiome functioning of the different bird species analyzed here likely influences their acquisition of antibiotic resistant bacteria. Penguins and avocets hunt small aquatic animals, while ducks filter water and sediments to trap plant and animal material. It is possible that ducks ingest large amounts of bacteria while dabbling. In addition, birds may have historical-evolutionary associations with particular microbial species, resulting in a distinct microbiome composition and functioning across avian taxonomic groups. Indeed, a metabarcoding study showed that bird taxonomy explained most of the compositional variation in the microbiome of birds (50). Our functional analyses also suggest that the different bird orders harbor microbial communities with distinct metabolisms. The microbiome of Anseriformes (ducks) expressed genes encoding significantly different metabolic pathways compared with other birds, while there was no clear distinction among Charadriiformes and Sphenisciformes (Fig. S5, Table S5). It is therefore plausible that the high resistance gene expression in ducks from the WWTP is influenced by their distinct microbiome, which in turn reflects their ecological niche and established host-microbe associations. In this scenario, bird traits modulate (amplifying or diminishing) the human impact on the spread of resistance genes.

Migratory birds are of particular concern as they might spread antibiotic resistance across large geographic distances in the same way that they disperse pathogens (9, 11, 51, 52). There are significant differences in gut microbiomes of migratory and resident red-necked stints (*Calidris ruficolis*) and curlew sandpipers (*Calidris furringea*), although these differences may be temporary (53, 54). Ruddy turnstones have a remarkable migration habit, travelling between breeding areas in Siberia to non-breeding sites in Australia via East-Asia, potentially acquiring and distributing resistant bacteria along the way. The turnstones analyzed here carried resistance against several antibiotic classes, but the diversity of genes within those classes was much smaller than in birds at the WWTP (Fig. 1). *Anas* ducks travel hundreds of kilometers within Australia (55). It is possible that ducks from the Innamincka reserve have been in sites of high human impact previously, resulting in the higher load of resistance genes when compared with other birds from remote areas. It is also plausible that ducks acquire resistant bacteria due to their feeding behavior and the composition of their gut microbiome.

Despite their isolation, we found genes conferring resistance against four antibiotic classes in penguins from Antarctica. Previous studies of antibiotic resistance in penguins have produced contradictory results. In one, various tetracycline resistant bacteria were isolated from the cloaca of penguins (56), while in another high levels of resistance against multiple antibiotics were detected in penguin droppings (57). However, other studies have reported that antibiotic resistant bacteria are rare in these animals (24, 58, 59). It is possible that penguins acquire resistance genes from migratory fish and other prey or animals with which they interact. As antibiotics are naturally produced by bacteria, it is also possible that the resistance genes observed in the penguin microbiome occur in the environment regardless of human influence. The possibility of some cross-library and/or environmental contamination cannot be completely excluded. Nevertheless, the *bona fide* influence of human activity is supported by the larger number of resistance genes adjacent to the more populated O’Higgins Base compared with the much smaller González Videla Base. Additionally, previous research shows higher antibiotic resistance levels near research facilities compared to more pristine sites in Antarctica (24, 57). Increasing research activities, tourism and limited sewage treatment (60) are therefore the most likely explanation for the presence of antibiotic resistance in Antarctic penguins.

The bird microbiome expressed resistance against nine classes of antibiotics, even though we putatively enriched libraries with resistant bacterial strains using only two classes of antibiotics in the collection media (aminoglycoside and β-lactams, see Materials and Methods). Acquired (horizontally transferred) resistance genes can be constitutively expressed, in which case the presence of their transcripts is expected even without antibiotic exposure. It is also possible that these resistance genes were acting against antibiotics present in the environment and/or that these genes are co-transmitted with others that have functions in addition to antibiotic resistance (e.g. metal resistance, 61). Variables related to the ecology and geographic distribution of the different bird species could also play a role, although the results based on individual collection sites and bird taxonomic group show that these variables are unlikely to change the conclusion that birds from the WWTP carry the highest diversity and abundance of resistance genes. Meta-transcriptomic studies necessarily rely on reference databases, which limits the discovery of novel resistance genes (30), and the database used here (ResFinder, 62) does not include resistance that arises through *de novo* mutation in the bacterial genome (which would increase the detection of false positives). Therefore, although we were limited to assessing acquired resistance genes, these genes residing on mobile elements pose greater public health risk as they can be transferred easily between bacteria (63). Potentially unequal RNA yields across libraries represent an additional caveat. We observed no correlation between library size or microbial mRNA reads and the diversity or abundance of resistance genes, indicating that the higher number of resistance genes observed in the WWTP does not result from unequal sequencing effort, or from failing to extract and sequence microbes from other sites (Fig. S6). Considering our rather conservative analyses (see Materials and Methods), it is possible that we underestimate the presence of some resistance genes that were not expressed or were expressed at low abundance.

In sum, we show that ducks feeding on wastewater are particularly prone to harbor bacteria with transcriptionally active antibiotic resistance genes. Ecological and functional traits are likely intertwined in explaining the higher propensity of ducks to carry antibiotic resistance genes. This study also contributes to the increasing literature reporting widespread antibiotic resistance in birds, even in isolated areas like the Australian outback and Antarctica. For antibiotic resistant bacteria, aquatic systems are major traffic routes between wildlife and humans (8, 64). The resistance genes acquired by birds can be re-introduced in the environment, possibly in other water systems (e.g. by migrating ducks) and might re-infect humans directly via contact with contaminated water, or indirectly by the introduction of these genes into the food chain (64). Investigating the mechanisms that sustain the persistence and cycling of resistance genes in wild populations despite the metabolic burden that these gene impose is a logical next step towards tackling antibiotic resistance.

## Materials and Methods

### Sampling

Samples were collected as part of long-term avian influenza virus surveillance studies (65-70). Ethics approvals, bird capture methods and sample handling are reported in the Supplementary Materials. In short, cloacal and oropharyngeal swabs were collected using a sterile-tipped applicator and placed in viral transport media (VTM, Brain-heart infusion broth containing 2 × 106IU/l penicillin, 0.2mg/ml streptomycin, 0.5mg/ml gentamicin, 500U/ml amphotericin B, Sigma). VTM is a standard buffer used in avian influenza surveys and has the advantage of killing a portion of non-resistant bacterial strains. This step enriches meta-transcriptomes libraries with antibiotic resistant bacteria and, consequently, increases the sensitivity of the antibiotic resistance survey. As all samples were stored in VTM, there is no reason to believe that this step would affect the abundance comparisons among libraries, and thus, it is unlikely to bias the results. All birds in this study were apparently healthy, with the exception of one library constructed from dead and dying shelducks with symptoms of Newcastle Disease. Samples were assayed for avian influenza virus as previously described (66). Samples were collected at sites with different levels of anthropogenic impact (Supplementary Materials). Birds sampled at the WWTP were found in lagoons composed of partially treated water (the final stage of wastewater treatment).

### RNA-sequencing and data processing

RNA isolation procedures are detailed in the Supplementary Materials. Libraries were composed of 10 conspecific bird samples pooled at equal concentrations. Paired-end sequencing (100bp) was performed on a HiSeq2500 platform and the number of reads obtained are reported in Table S1. Low quality reads, adapters, host reads and ribosomal RNA were filtered out from the data set (Supplementary Materials).

### Resistance genes characterization

The ResFinder reference database (62) was used in conjunction with the KMA program (71) (downloaded in December 2017) to identify resistance genes in the meta-transcriptomic data set. The ResFinder database currently contains 2255 resistance genes compiled from published manuscripts and existing databases. KMA was preferred over other alignment tools because it performs well in aligning short reads against highly redundant databases and is able to resolve non-unique read matches by assessing and statistically testing global alignment scores. To minimize the risk of false-positives and increase the minimum mapping length allowed, only genes with a mapping coverage greater than 20% were considered in the analyses, all of with had an alignment *p*-value << 0.05. The average length of the resistance genes observed was 944bp – a gene with this length was only considered in the downstream analyses if query reads overlapped by at least 189 bp (20% coverage). This approach is highly conservative because it uses an aligner that yields a minimal number of false positives (71), does not include housekeeping genes (which would increase the occurrence of false positives), and defines resistance genes based on gene fragments (at least 20% of the genes) rather than individual reads. The gene fragments analyzed here are longer than the ones obtained via qPCR (generally 100bp amplicons), which are widely used in AMR assessments of environmental samples and in diagnostic laboratories. One gene (*bla*_TEM-116_) was observed in all libraries but was removed from the analyses due to its potential contaminant nature (72). It is possible that the data set contains other laboratory contaminants, but the fact that one of the libraries contained only one resistance gene, and that no other gene (except for *bla*_TEM-116_) was found in all libraries, suggests that contamination is unlikely. Genes conferring resistance to Macrolide, Lincosamide and Streptogramin B were considered as one antibiotic class (MLS). Absolute read abundances were estimated based on a stably expressed host gene and normalized for gene length (Supplementary Materials). The number of antibiotic classes to which resistance was found, the diversity (i.e. number of genes) and the abundance of resistance genes in each library were classified into two bins (‘WWTP’ and ‘Other’, Fig. 2). Differences between WWTP and other sites were tested with a Kruskal-Wallis test using the native *stats* R package (R Core Team 73). The higher diversity of resistance genes in libraries from the WWTP was validated with a PCR-based approach targeting resistance genes in individual birds from two libraries (n=20, Supplementary Materials).

### Functional profiling

The microorganism-based functional profile was inferred with HUMAnN2 (74) (http://huttenhower.sph.harvard.edu/humann2), using the UniRef90 protein database as reference (75). Community-level differences in expression of pathways between sites and bird orders was visually assessed with Principal Coordinate Analysis based on an Euclidean distance matrix with the *ape* R package (76) and further investigated with Random Forest analysis, using 1000 trees, with the *randomForest* R package (77).

## Acknowledgments

The sampling was supported by NIAID (HHSN266200700010C), ARC discovery grants (DP 130101935 and DP160102146) and the Instituto Chileno Antártico INACH (T 12-13 and T 27-10). The Melbourne WHO Collaborating Centre for Reference and Research on Influenza is supported by the Australian Department of Health. ECH is funded by an ARC Australian Laureate Fellowship (FL170100022). TCS is a Sydney Medical Foundation Fellow whose work is also supported by the Sydney Medical School Foundation. JRI is supported by a NHMRC Practitioner Fellowship (GNT1104232). We thank the High Performance Computing team at Sydney University, the genomic facilities at the Westmead Institute for Medical Research, the Centre for Integrative Ecology at Deakin University, the Victorian Wader Study Group, and the logistic support from Melbourne Water, Innamincka Station and Innamincka Regional Reserve. We also thank Sebastiaan van Hal, Sally Partridge, Ali Khalid, Philip Clausen, Simeon Lisovski and Marta Ferenczi for their support.

## Authors’ contributions

VRM, MW and ECH designed the research. MW, DG-A and MK collected samples. MW carried out DNA/RNA isolation and meta-transcriptome library preparation. VRM and J-SE performed PCRs. VRM performed the data analyses. ACH, DG-A, MK, TCS and ECH contributed with reagents and/or funds for research. All authors contributed to interpreting data and manuscript writing. All authors gave final approval for publication.

## References

1. Van Boeckel TP, et al. (2017) Reducing antimicrobial use in food animals. Science 357(6358):1350–1352.

2. Klein EY, et al. (2018) Global increase and geographic convergence in antibiotic consumption between 2000 and 2015. Proc Natl Acad Sci U S A 115(15):E3463–E3470.

3. O’Neill J (2016) Tackling drug-resistant infections globally: final report and recommendations. (amr-review.org).

4. D’Costa VM, et al. (2011) Antibiotic resistance is ancient. Nature 477(7365):457–461.

5. Bhullar K, et al. (2012) Antibiotic resistance is prevalent in an isolated cave microbiome. PLoS One 7(4):e34953.

6. Harmer CJ & Hall RM (2015) The A to Z of A/C plasmids. Plasmid 80:63–82.

7. Crofts TS, Gasparrini AJ, & Dantas G (2017) Next-generation approaches to understand and combat the antibiotic resistome. Nat. Rev. Microbiol. 15(7):422–434.

8. Vittecoq M, et al. (2016) Antimicrobial resistance in wildlife. J. Appl. Ecol. 53(2):519–529.

9. Surette MD & Wright GD (2017) Lessons from the environmental antibiotic resistome. Annu. Rev. Microbiol. 71:309–329.

10. Stedt J, et al. (2014) Antibiotic resistance patterns in *Escherichia coli* from gulls in nine European countries. Infect Ecol Epidemiol 4.

11. Bauer S & Hoye BJ (2014) Migratory animals couple biodiversity and ecosystem functioning worldwide. Science 344(6179):1242552.

12. Atterby C, et al. (2016) Increased prevalence of antibiotic-resistant. *E. coli* in gulls sampled in Southcentral Alaska is associated with urban environments. Infect Ecol Epidemiol 6:32334.

13. Stedt J, et al. (2015) Carriage of CTX-M type extended spectrum beta-lactamases (ESBLs) in gulls across Europe. Acta Vet Scand 57:74.

14. Hasan B, et al. (2012) Antimicrobial drug-resistant. *Escherichia coli* in wild birds and free-range poultry, Bangladesh. Emerg Infect Dis 18(12):2055–2058.

15. Atterby C, et al. (2017) ESBL-producing *Escherichia coli* in Swedish gulls-A case of environmental pollution from humans. PLoS One 12(12):e0190380.

16. Hernandez J, et al. (2013) Characterization and comparison of extended-spectrum beta-lactamase (ESBL) resistance genotypes and population structure of *Escherichia coli* isolated from Franklin’s gulls (*Leucophaeus pipixcan*) and humans in Chile. PLoS One 8(9):e76150.

17. Martiny AC, Martiny JB, Weihe C, Field A, & Ellis JC (2011) Functional metagenomics reveals previously unrecognized diversity of antibiotic resistance genes in gulls. Front. Microbiol. 2:238.

18. Dolejska M, Cizek A, & Literak I (2007) High prevalence of antimicrobial-resistant genes and integrons in *Escherichia coli* isolates from Black-headed Gulls in the Czech Republic. J. Appl. Microbiol. 103(1):11–19.

19. Veldman K, van Tulden P, Kant A, Testerink J, & Mevius D (2013) Characteristics of cefotaxime-resistant *Escherichia coli* from wild birds in the Netherlands. Appl. Environ. Microbiol. 79(24):7556–7561.

20. Ahlstrom CA, et al. (2018) Acquisition and dissemination of cephalosporin-resistant *E. coli* in migratory birds sampled at an Alaska landfill as inferred through genomic analysis. Sci. Rep. 8(1):7361.

21. Chu BTT, et al. (2017) Metagenomic analysis reveals the impact of wastewater treatment plants on the dispersal of microorganisms and genes in aquatic sediments. Appl. Environ. Microbiol.

22. Pehrsson EC, et al. (2016) Interconnected microbiomes and resistomes in low-income human habitats. Nature 533(7602):212–216.

23. Zhang XX, Zhang T, & Fang HH (2009) Antibiotic resistance genes in water environment. Appl. Microbiol. Biotechnol. 82(3):397–414.

24. Rabbia V, et al. (2016) Antibiotic resistance in *Escherichia coli* strains isolated from Antarctic bird feces, water from inside a wastewater treatment plant, and seawater samples collected in the Antarctic Treaty area. Polar Science 10(2):123–131.

25. Rodriguez-Mozaz S, et al. (2015) Occurrence of antibiotics and antibiotic resistance genes in hospital and urban wastewaters and their impact on the receiving river. Water Res. 69:234–242.

26. Mohammadali M & Davies J (2017) Antimicrobial Resistance Genes and Wastewater Treatment. Antimicrobial Resistance in Wastewater Treatment Processes, eds Keen PL & Fugère R), pp 1–13.

27. Zhu YG, et al. (2017) Continental-scale pollution of estuaries with antibiotic resistance genes. Nat Microbiol 2:16270.

28. Zhao Y, et al. (2017) Feed additives shift gut microbiota and enrich antibiotic resistance in swine gut. Sci. Total Environ.

29. Zhu YG, et al. (2013) Diverse and abundant antibiotic resistance genes in Chinese swine farms. Proc Natl Acad Sci U S A 110(9):3435–3440.

30. Bengtsson-Palme J, Larsson DGJ, & Kristiansson E (2017) Using metagenomics to investigate human and environmental resistomes. J. Antimicrob. Chemother. 72(10):2690–2703.

31. Su JQ, et al. (2017) Metagenomics of urban sewage identifies an extensively shared antibiotic resistome in China. Microbiome 5(1):84.

32. Munk P, et al. (2018) Abundance and diversity of the faecal resistome in slaughter pigs and broilers in nine European countries. Nat Microbiol 3(8):898–908.

33. Mira A, Ochman H, & Moran NA (2001) Deletional bias and the evolution of bacterial genomes. Trends Genet. 17:589–596.

34. Hessen DO, Jeyasingh PD, Neiman M, & Weider LJ (2010) Genome streamlining and the elemental costs of growth. Trends Ecol. Evol. 25(2):75–80.

35. Marcelino VR, Cremen MC, Jackson CJ, Larkum AWD, & Verbruggen H (2016) Evolutionary dynamics of chloroplast genomes in low light: a case study of the endolithic green alga *Ostreobium quekettii*. Genome Biol. Evol. 8(9):2939–2951.

36. Wolf YI & Koonin EV (2013) Genome reduction as the dominant mode of evolution. Bioessays 35(9):829–837.

37. Versluis D, et al. (2015) Mining microbial metatranscriptomes for expression of antibiotic resistance genes under natural conditions. Sci. Rep. 5:11981.

38. Rowe WPM, et al. (2017) Overexpression of antibiotic resistance genes in hospital effluents over time. J. Antimicrob. Chemother. 72(6):1617–1623.

39. Paliy O & Shankar V (2016) Application of multivariate statistical techniques in microbial ecology. Mol. Ecol. 25(5):1032–1057.

40. Ganz HH, et al. (2017) Community-level differences in the microbiome of healthy wild mallards and those infected by influenza A viruses. mSystems 2(1).

41. Walsh C (2000) Molecular mechanisms that confer antibacterial drug resistance. Nature 406(6797):775–781.

42. D’Andrea MM, Arena F, Pallecchi L, & Rossolini GM (2013) CTX-M-type beta-lactamases: a successful story of antibiotic resistance. Int. J. Med. Microbiol. 303(6-7):305–317.

43. Falagas ME, Grammatikos AP, & Michalopoulos A (2008) Potential of old-generation antibiotics to address current need for new antibiotics. Expert Rev Anti Infect Ther 6(5):593–600.

44. Arcangioli MA, Leroy-Setrin S, Martel JL, & Chaslus-Dancla E (1999) A new chloramphenicol and florfenicol resistance gene flanked by two integron structures in Salmonella typhimurium DT104. FEMS Microbiol. Lett. 174(2):327–332.

45. Cloeckaert A, et al. (2000) Plasmid-mediated florfenicol resistance encoded by the floR gene in Escherichia coli isolated from cattle. Antimicrob. Agents Chemother. 44(10):2858–2860.

46. Berendsen BJA, et al. (2018) The persistence of a broad range of antibiotics during calve, pig and broiler manure storage. Chemosphere 204:267–276.

47. Poirel L, Cattoir V, & Nordmann P (2012) Plasmid-mediated quinolone resistance; interactions between human, animal, and environmental ecologies. Front. Microbiol. 3:24.

48. Arnott A, et al. (2018) Multidrug-Resistan. *Salmonella enterica* 4,[5],12:i:-Sequence Type 34, New South Wales, Australia, 2016-2017. Emerg Infect Dis 24(4):751–753.

49. Robicsek A, et al. (2006) Fluoroquinolone-modifying enzyme: a new adaptation of a common aminoglycoside acetyltransferase. Nat. Med. 12(1):83–88.

50. Hird SM, Sanchez C, Carstens BC, & Brumfield RT (2015) Comparative Gut Microbiota of 59 Neotropical Bird Species. Front. Microbiol. 6:1403.

51. Altizer S, Bartel R, & Han BA (2011) Animal migration and infectious disease risk. Science 331(6015):296–302.

52. Fourment M, Darling AE, & Holmes EC (2017) The impact of migratory flyways on the spread of avian influenza virus in North America. BMC Evol. Biol. 17(1):118.

53. Risely A, Waite DW, Ujvari B, Hoye BJ, & Klaassen M (2017) Active migration is associated with specific and consistent changes to gut microbiota in Calidris shorebirds. J. Anim. Ecol.

54. Risely A, Waite D, Ujvari B, Klaassen M, & Hoye B (2017) Gut microbiota of a long-distance migrant demonstrates resistance against environmental microbe incursions. Mol. Ecol. 26(20):5842–5854.

55. Roshier D, Asmus M, & Klaassen M (2008) What drives long-distance movements in the nomadic Grey Teal *Anas gracilis* in Australia. Ibis 150(3):474–484.

56. Rahman MH, Sakamoto KQ, Kitamura S-I, Nonaka L, & Suzuki S (2015) Diversity of tetracycline-resistant bacteria and resistance gene tet(M) in fecal microbial community of Adélie penguin in Antarctica. Polar Biol. 38(10):1775–1781.

57. Miller RV, Gammon K, & Day MJ (2009) Antibiotic resistance among bacteria isolated from seawater and penguin fecal samples collected near Palmer Station, Antarctica. Can J Microbiol 55(1):37–45.

58. Bonnedahl J, et al. (2008) Antibiotic susceptibility of faecal bacteria in Antarctic penguins. Polar Biol. 31(6):759–763.

59. Hernandez J, et al. (2012) Human-associated extended-spectrum beta-lactamase in the Antarctic. Appl. Environ. Microbiol. 78(6):2056–2058.

60. Gröndahl F, Sidenmark J, & Thomsen A (2016) Survey of waste water disposal practices at Antarctic research stations. Polar Res. 28(2):298–306.

61. Baker-Austin C, Wright MS, Stepanauskas R, & McArthur JV (2006) Co-selection of antibiotic and metal resistance. Trends Microbiol. 14(4):176–182.

62. Zankari E, et al. (2012) Identification of acquired antimicrobial resistance genes. J. Antimicrob. Chemother. 67(11):2640–2644.

63. Martinez JL, Coque TM, & Baquero F (2015) What is a resistance gene? Ranking risk in resistomes. Nat. Rev. Microbiol. 13(2):116–123.

64. Taylor NG, Verner-Jeffreys DW, & Baker-Austin C (2011) Aquatic systems: maintaining, mixing and mobilising antimicrobial resistance. Trends Ecol. Evol. 26(6):278–284.

65. Ferenczi M (2016) Avian influenza virus dynamics in Australian wild birds. PhD. (Deakin Univeristy).

66. Ferenczi M, et al. (2016) Avian influenza infection dynamics under variable climatic conditions, viral prevalence is rainfall driven in waterfowl from temperate, south-east Australia. Vet Res 47:23.

67. Hurt AC, et al. (2014) Detection of evolutionarily distinct avian influenza a viruses in antarctica. MBio 5(3):e01098–01014.

68. Hurt AC, et al. (2016) Evidence for the introduction, reassortment, and persistence of diverse influenza A viruses in Antarctica. J. Virol. 90(21):9674–9682.

69. Grillo VL, et al. (2015) Avian influenza in Australia: a summary of 5 years of wild bird surveillance. Aust. Vet. J. 93(11):387–393.

70. González-Acuña D, et al. (2013) Health evaluation of wild gentoo penguins (*Pygoscelis papua*) in the Antarctic Peninsula. Polar Biol. 36(12):1749–1760.

71. Clausen P, Aarestrup FM, & Lund O (2018) Rapid and precise alignment of raw reads against redundant databases with KMA. BMC Bioinformatics 19(1):307.

72. Jacoby GA & Bush K (2016) The Curious Case of TEM-116. Antimicrob. Agents Chemother. 60(11):7000.

73. R Core Team (2013) R: A language and environment for statistical computing (Vienna, Austria.).

74. Abubucker S, et al. (2012) Metabolic reconstruction for metagenomic data and its application to the human microbiome. PLoS Comput Biol 8(6):e1002358.

75. Suzek BE, et al. (2015) UniRef clusters: a comprehensive and scalable alternative for improving sequence similarity searches. Bioinformatics 31(6):926–932.

76. Paradis E, Claude J, & Strimmer K (2004) APE: analyses of phylogenetics and evolution in R language. Bioinformatics 20:289–290.

77. Liaw A & Wiener M (2002) Classification and regression by randomForest. R News 2(3):18–22.

